# Simulations reveal high efficiency and confinement of a population suppression CRISPR toxin-antidote gene drive

**DOI:** 10.1101/2022.10.30.514459

**Authors:** Yutong Zhu, Jackson Champer

## Abstract

Though engineered gene drives hold great promise for spreading through and eventually suppressing populations of disease vectors or invasive species, complications such as resistance alleles and spatial population structure can prevent their success. Additionally, most forms of suppression drives, such as homing drives or driving Y chromosomes, will generally spread uncontrollably between populations with even small levels of migration. The previously proposed CRISPR-based toxin-antidote system called TADE suppression drive could potentially address the issue of confinement and resistance alleles. However, it is a relatively weak form of drive compared to homing drives, which might make it particularly vulnerable to spatial population structure. In this study, we investigate TADE suppression drive using individual-based simulations in continuous space. We find that the drive is actually more confined in continuous space than in panmictic populations, even in its most efficient form with a low cleavage rate in embryos from maternally deposited Cas9. Furthermore, the drive performed well in continuous space scenarios if the initial release requirements were met, suppressing the populations in a timely manner without being severely affected by chasing, a phenomenon in which wild-type individuals avoid the drive by recolonizing empty areas. At higher embryo cut rates, the drive loses its ability to propagate on its own, but a single, widespread release can often still induce rapid population collapse. Thus, if TADE suppression gene drives can be successfully constructed, they may play an important role in control of disease vectors and invasive species when stringent confinement to target populations is desired.

## Introduction

A gene drive allele can increase its frequency in a population^1–6^. These constructs have many possible mechanisms, and experimental demonstrations have been conducted in a wide variety of organisms, including yeast^7–10^, flies^11–28^, mosquitoes^29–33^, and mice^34^. Modification gene drives can carry a cargo or make other genetic changes for a variety of purposes such as to prevent vector-borne disease transmission, rescue an endangered species from disease, or even to support scientific studies. If a gene drive allele is designed to disrupt an essential gene without rescue, bias the sex ratio, or cause some other type of major fitness cost, it can potentially be used for population suppression as well. This can also support reduction of vector-borne diseases, but suppression drives can additionally be used to control invasive species and agricultural pests in a specific and environmentally friendly manner.

Most gene drives developed thus far are homing drives based on CRISPR/Cas9 nucleases together with guide RNAs (gRNAs), which can be used for modification or suppression and can increase in frequency very rapidly^1–6^. However, homing drives tend to be vulnerable to resistance alleles, mutations that cause mismatches with the drive’s gRNA(s) and thus block DNA cleavage and drive spread^24,35^. While it may be possible to mitigate the development of resistance alleles with improved drive designs^27,29^, homing drives also tend to spread if even a small number of drive-carrying individuals are present in a population. In some situations, this can be desirable, particularly for global elimination of diseases such as malaria or dengue. However, sociopolitical considerations or the need to control invasive species and agricultural pests in only target regions can make such uncontrolled spread undesirable, so we require development of methods that can confine a drive to a target population.

There are many examples of potentially confinable modification gene drives with a variety of mechanisms. These are confined via an “introduction threshold”, which can vary substantially between different types of drives. If introduced into a population at a frequency above the threshold, the drive will increase in frequency. If introduced below, the drive will tend to be removed from the population. Such introduction thresholds imply the existence of (lower) migration thresholds. In these settings, if the drive spreads in a target population, it could never reach significant frequency in a connected population if the migration level between the populations is below the threshold. Many confined drive systems have been demonstrated in fruit flies, such as the *Medea* toxin-antidote system^36^, a single locus underdominance TA system^15^, synthetic species underdominance^19^, and the CRISPR-based TARE/ClvR drives^16,17^. These latter systems target and disrupt an essential but haplosufficient gene with CRISPR (without being copied to the site of the DNA break), but the gene drive contains a functional copy of the target gene that is immune to cleavage. Thus, wild-type alleles are converted to disrupted alleles, which are removed from the population, increasing the relative frequency of the gene drive. Because multiplexed gRNAs can be used without the loss of efficiency seen in homing drives^21^, such CRISPR toxin-antidote drives are less vulnerable to resistance alleles^37^.

None of these forms of confined drive could support strong suppression. However, one similar CRISPR toxin-antidote design called “TADE (toxin-antidote dominant embryo) suppression” is capable of this^37,38^. Such a design has two important differences from TARE/ClvR. First, it targets a haplolethal gene instead of a haplosufficient but essential gene. This means that two copies of the target gene are required for viability, not just one. Second, the gene drive must disrupt a one-sex haplosufficient but essential gene (usually modeled as a female fertility gene). Such disruption can be from the presence of the drive inside the gene itself or from additional gRNAs added to the gene drive that target the gene (without rescue by the drive - an important distinction from the haplolethal target gene for which the drive provides rescue). As the drive increases in frequency due to its rescue element, it is this disruption of the one-sex haplosufficient but essential gene that induces population suppression. Like TARE drive, TADE suppression drive lacks an introduction threshold in idealized form, but the threshold immediately appears and increases steeply for any reduction in drive efficiency or embryo cut rate^37,38^. Though not yet constructed, TADE suppression drive could feasibly be developed using off-the-shelf components from other gene drives targeting haplolethal^27^ and female fertility^20^ genes. Despite the greater difficulty of engineering constructs with haplolethal targets, is thus a feasible and promising candidate for confined suppression drive.

However, TADE suppression drive has thus far only been modeled in simple panmictic population. Previous studies have shown that drive properties such as confinement can substantially vary in spatially explicit populations for frequency-dependent, confined drive systems^39–41^. Suppression gene drives have additional complexities in spatially explicit populations^42–50^. Perhaps the most prominent of these is the “chasing” effect, in which some wild-type individuals avoid suppression by the gene drive and reach areas of empty space where the population was previously suppressed by the drive. With reduced competition, the wild-type individuals are then able to expand quickly into this empty space, recolonizing the area before the gene drive returns (“chasing” the wild-type population through the model arena) to again suppress the local population and continue the cycle. It is possible that with its slower rate of spread, a confined, frequency-dependent gene drive such as TADE suppression drive would be even more vulnerable to chasing than previously tested suppression drives based on homing or X-chromosome shredding^42,47^.

In this study, we use individual-based simulations to assess the performance of TADE suppression drive in continuous space. We find that it is substantially more confined in this setting than in panmictic populations. However, if the drive does establish, it is often able to successfully suppress populations with similar rates of success as other suppression drives. Less effective versions of TADE suppression drive that are unable to establish after a single, local release can also be used to effectively suppress populations with a strategy based on a single, widespread release of drive-carrying individuals. All these characteristics make TADE suppression drives potentially useful tools for elimination of target pest populations.

## Methods

### TADE Suppression drive

All TADE drives target and disrupt a haplolethal gene with a nuclease (generally using CRISPR/Cas9 together with multiple gRNAs to ensure that the target gene is rendered nonfunctional - see Figure 1)^37,38^. Two functional copies of the haplolethal target gene are required for viability. These could be provided by wild-type alleles or by drive alleles, which contain a recoded copy of the target gene that is immune to drive-induced cleavage. TADE suppression drives must also disrupt a haplosufficient but essential female fertility gene, so that only females with at least one wild-type copy of this gene can reproduce.

**Figure 1.**
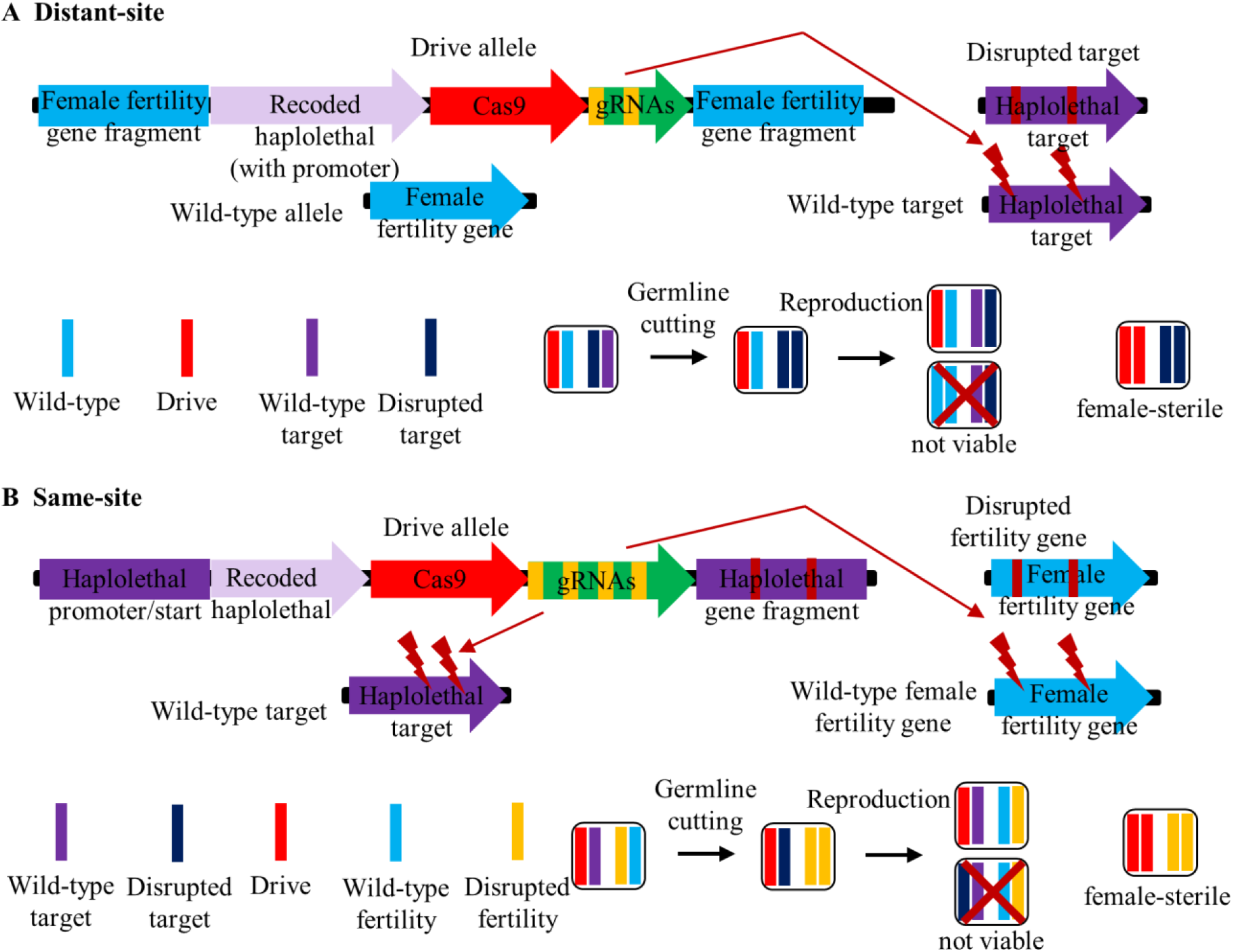
TADE suppression drive components and mechanism. (**A**) A distant-site TADE suppression drive is placed inside a female fertility gene, disrupting it. It targets a distant haplolethal gene with Cas9 and gRNAs, while providing a full rescue gene, including a promoter. (**B**) A same-site TADE suppression drive is placed inside the haplolethal target gene, providing rescue with some additional recoded sequence. It also targets a distant female fertility gene with Cas9 and gRNAs.

In “distant site” TADE suppression drive, the drive is located inside the female fertility gene, disrupting it (Figure 1A). Thus, drive homozygous females are sterile. The drive contains a complete recoded copy of the haplolethal target gene, including promoter elements. In “same-site” TADE suppression drive, the drive is located in the haplolethal target gene, inserted at a different site than the gRNA target sites^37,38^ (Figure 1B). Thus, the rescue copy of the drive uses native promoter elements. This drive contains additional gRNAs that target and disrupt the female fertility gene. Each of these configurations has potential advantages, though both have nearly identical performance when cut rates are high^37,38^. The same-site method allows for a greater chance of successful rescue of the target gene, a requirement for drive function, as well as a smaller rescue element that need not include gene regulatory elements. The distant-site method has better performance when cut rates are low and does not require separate gRNAs targeting the female fertility gene.

Here, we explicitly model distant-site TADE suppression drives, though we use high germline cuts rates, making our results fully applicable to same-site drives. Any genotype that does not have at least two alleles with functional haplolethal target gene (wild-type and drive alleles) is considered to be nonviable. Further, any female drive homozygotes are considered to be sterile due to lack of any functional female fertility genes. In the germline, wild-type target alleles are converted to disrupted alleles at a rate equal to the germline cut rate if any drive alleles are present. These alleles are then passed on to offspring. Our default germline cut rate is 99% (which has been achieved in several experimental studies in flies and *Anopheles* mosquitoes), with an ideal cut rate of 100%. Maternal Cas9/gRNA deposition can also convert wild-type alleles in new embryos to disrupted alleles if the female parent has a drive allele. The rate that this occurs is equal to the embryo cut rate. The default value for this parameter is 5%, representing a high quality but imperfect drive, with an ideal rate of 0%.

### Simulation model

This model is based on the previous studies^42,47^. We simulated sexually reproducing diploids with SLiM software (version 3.7)^51^. We tracked the position of each individual in a 1×1 (unitless) arena. Our model has discrete, non-overlapping generations.

The simulations are initialized by randomly placing 50,000 wild-type individuals in the arena. We run the simulation for 10 generation to allow the population to equilibrate, and we then release gene drive heterozygotes at “generation zero”. Unless otherwise specified, 33000 gene drive heterozygotes are released over a radius of 0.38 (representing ∼59% of individuals in the release area - these values were calibrated to allow the drive with default parameters to solidly establish in the release area, see Figure 3A). We end the simulation after 1000 generations or when the population or drive allele is eliminated.

We define fitness costs for drive homozygotes relative to wild-type individuals, with multiplicative fitness for each drive allele. The default value for drive homozygote fitness is 0.95 (wild-type fitness is 1), and the fitness cost of heterozygotes is the square root of this value. Fitness affects male mating success and female fecundity (see below).

### Reproduction in the simulations

Each fertile female will randomly select a male within the “migration rate” radius (which has a default of 0.04). She will then mate with this male with a probability equal to the male’s fitness. If she does not mate with this male, she will select another male to attempt to mate (with a maximum of 20 attempts). Females will not reproduce if they do not mate.

Female fecundity is affected by population density in a 0.01 radius around the female (*ρ*_*i*_), a carrying density constant (*ρ*, equal to population carrying capacity/total area), the fitness of the female (*f)* and population growth rate in low density (*β*). Specifically, the female fecundity is *ω*_*i*_ = *βf*/[(*β*-1) (*ρ*_*i*_/*ρ*)+1]. The number of offspring is determined by a random draw from a binomial distribution, with 50 chances to generate an offspring and a success probability for each chance equal to *ω*_*i*_/25.

Offspring are displaced from their mother in a random direction by a distance drawn from a normal distribution with a standard deviation equal to the migration rate. This produces an average displacement of 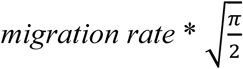. If an individual is placed outside the arena, its position is redrawn until it is placed in the arena.

### Data generation

Simulations were run on the High-Performance Computing Platform of the Center for Life Science at Peking University. Python was used to process data and prepare the figures. All SLiM models and scripts are available on Github (https://github.com/jchamper/ChamperLab/tree/main/TADE-Suppression-Modeling).

## Results

### Confinement of TADE suppression drive in spatially continuous populations

TADE drive works by targeting and disrupting a haplolethal gene, where two functional copies are needed for survival. The drive provides a recoded copy of the target gene, but it can still be eliminated from the population if a single drive allele is present without any functional wild-type target alleles. This occurs less often in an ideal TADE drive that has only germline cutting, usually requires mating between two different drive carriers. However, the drive can be lost in the progeny of female drive carriers when cleavage occurs in the embryo. In panmictic populations, this results in a heterozygote introduction threshold appearing and increasing to 61% for TADE suppression drive as the embryo cut rate increases to 100%^37,38^. This threshold can influence drive confinement and is relevant for models of two demes connected by migration.

However, the nature of confinement is different in spatially continuous models. Here, the drive can be established in one area, and then potentially advance directly into an adjacent area within the same region (Figure 2). It could also fail to persist, retreating in the face of a wild-type wave of advance. For modification drives, an introduction threshold of 50% corresponds approximately to the boundary between drive advance and retreat, though this can be slightly lower in drives with fitness costs^41^. However, for suppression drives, it could significantly change. This is because rather than advancing from a solid block of drive individuals, most space behind the advancing drive wave will consist of empty space due to the suppression effect, reducing the number of drive individuals that can advance forward by migration. This can influence a drive’s rate of advance^52^. To assess the ability of a TADE suppression to advance, we converted individuals in the left 25% of a square arena into drive heterozygotes. We then tracked the ability of the drive to reach the right portion of the arena, which indicated that the drive was able to form a wave of advance. With ideal parameters, TADE suppression drive was only able to reliably advance when the embryo cut rate was no more than 17% (Figure 2), which corresponds to a drive introduction threshold of 24%^38^, well below 50% and consistent with the notion that suppression drives may be substantially more confined in continuous space than in linked deme scenarios. For the drive with default parameters representing small imperfections, the drive could only reliably advance if the embryo cut rate was no more than 10%, a substantial reduction. TADE suppression drives must therefore have both low fitness costs and low embryo cuts rates to be able to effectively spread in spatially continuous scenarios.

**Figure 2.**
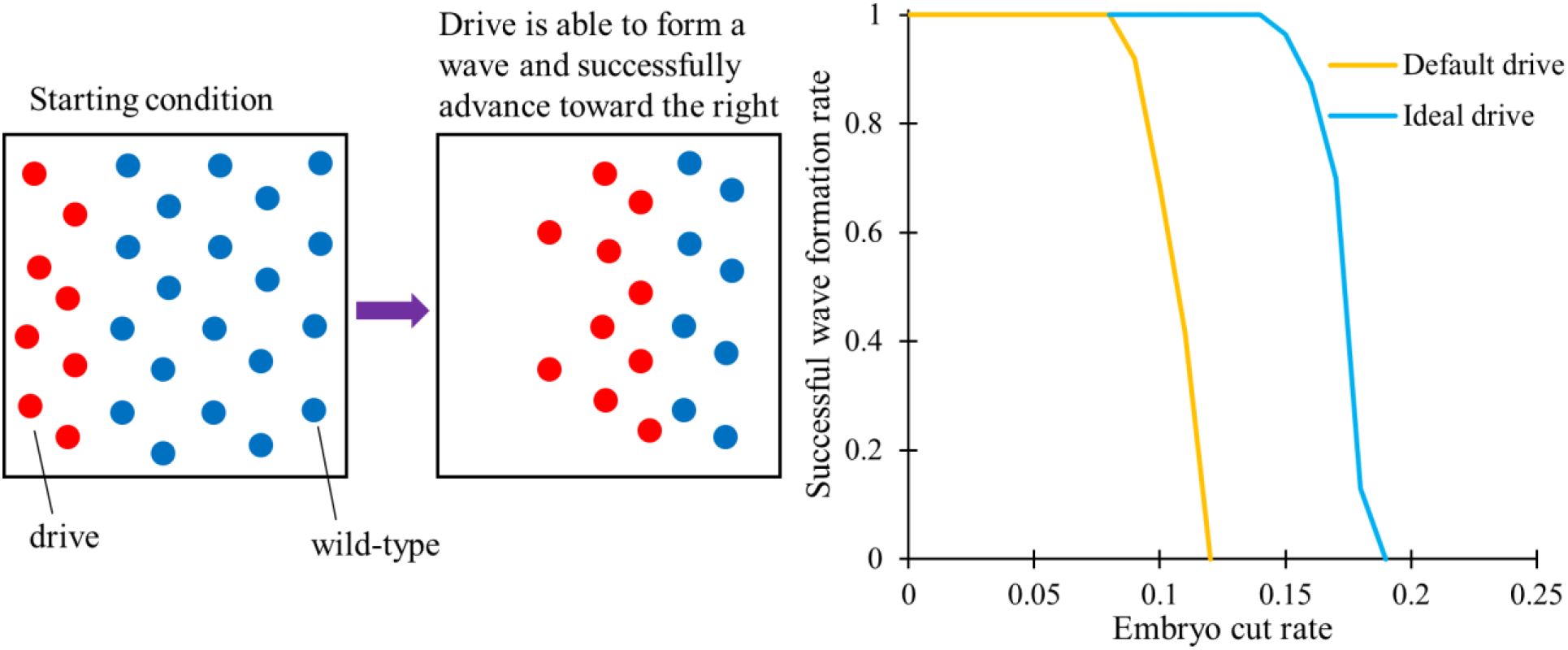
Effect of embryo cut rate on drive confinement. Individuals in the left side of a 1×1 arena with a population size of 50,000 were converted to drive heterozygotes out to a distance of 0.25. The simulation was allowed to run for several generations to measure if the drive could achieve high frequency on the right side of the arena. If so, this means that the drive could successfully form a wave of advance and could thus be expected to expand its range in spatial scenarios if it could initially establish. The right panel shows the fraction of simulations (out of 20 replicates per point) in which the drive was able to advance in space for both ideal and default performance parameters based on varying embryo cut rate.

Unlike homing drives and other zero-threshold systems that can quickly spread from small release sizes, frequency-dependent drives usually require at least a modest initial frequency to spread at an appreciable rate. Drives with thresholds must be released above their threshold to spread at all, and in spatially continuous scenarios, the area in which the drive is released must be sufficiently large to avoid being swamped by surrounding wild-type individuals^41^. This can result in drive elimination even if the drive is able to advance when it is not at a geometrical disadvantage. To assess the ability of TADE suppression drives to invade in a population after an initial release, we varied the release radius and release fraction in the area of release. In general, the drive with default parameters was only successful (avoiding drive loss) if the release radius was at least 0.3, with the release fraction being 25-50% (Figure 3A), despite this drive only having a panmictic introduction threshold of somewhat over 10%. An ideal TADE suppression drive with zero panmictic introduction threshold required a substantially lower introduction size to avoid drive loss, which only occurred due to stochastic factors at very low release sizes (Figure S1A). Thus, while an ideal TADE suppression drive can be easily introduced to a population with a modest release size, a substantially larger introduction size is required for imperfect versions.

**Figure 3.**
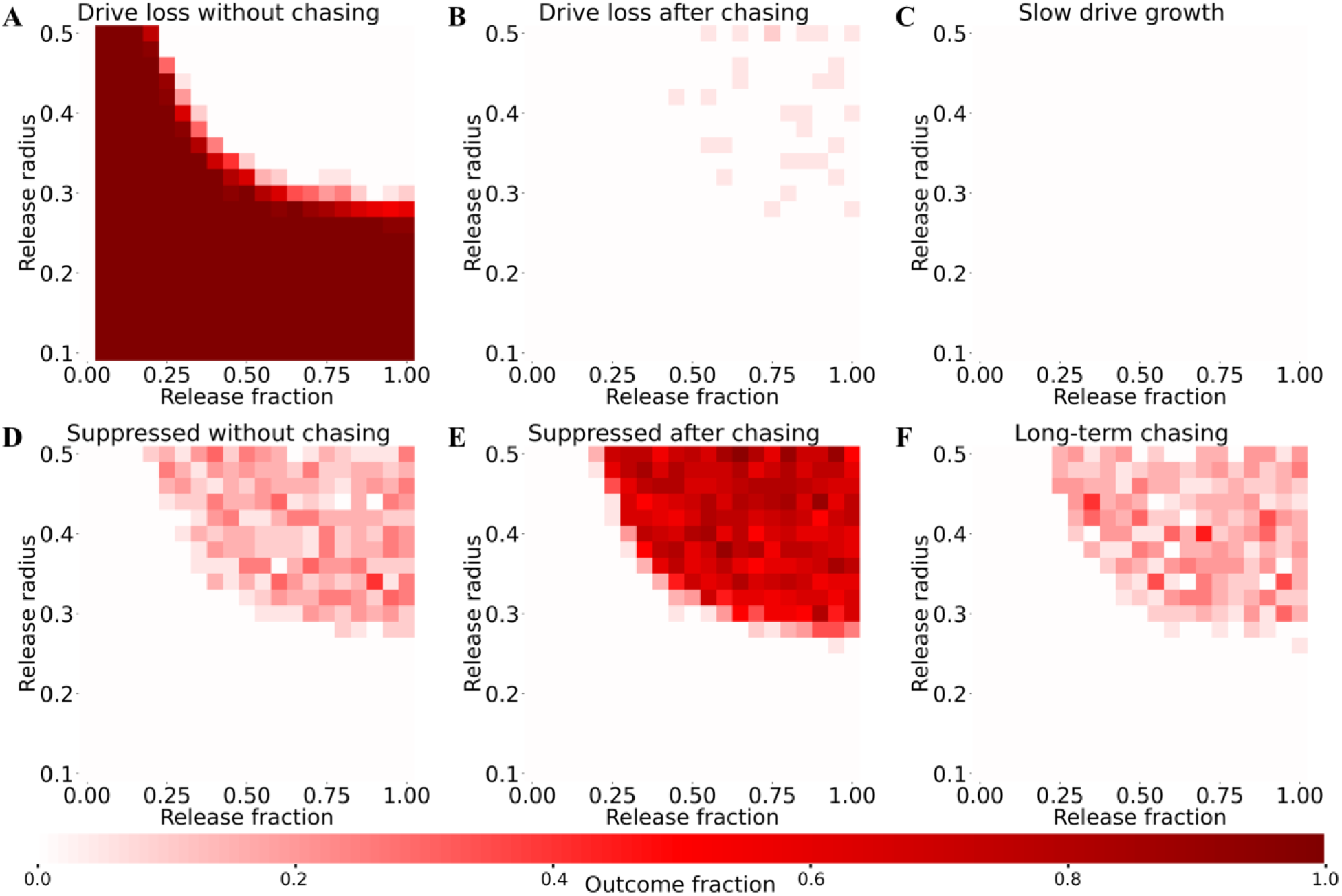
Outcomes with varying release parameters. Drive heterozygotes with default parameters were released into the middle of a spatial population of 50,000 individuals. The release radius and the fraction of drive individuals released (proportional to the average population in the release area) were varied. Outcome rates are displayed for (**A**) drive loss without chasing (this occurred due to inability of the drive to establish), (**B**) drive loss after a period of chasing, (**C**) simulations in which the drive remained in the population, but spread too slowly to achieve another type of outcome (this did not occur in these simulations, but see other data sets), (**D**) suppression without chasing, (**E**) suppression after a period of chasing, and (**F**) simulations in which chasing was still occurring after 1000 generations had elapsed. 20 simulations were assessed for each point in the parameter space.

### Drive outcomes in continuous space

Previous studies of suppression drive in continuous space have indicated that in many cases, suppression could be thwarted by the “chasing” phenomenon^42,47–50,53^. When this occurs, wild-type individuals escape into areas where the population was previously suppressed by the drive. With reduced competition, the wild-type individuals enjoy high reproductive success in these areas, which results in a rapid increase in the wild-type population. The drive is still present and can continue to locally suppress the population, but final suppression is delayed, perhaps indefinitely. For frequency-dependent TADE suppression drive, chasing could be more detrimental to the drive than for zero-threshold systems. This is because a rebounding wild-type population could reduce the drive to below its threshold level, potentially resulting in drive loss after a period of chasing. However, this was not observed to occur at high rates (Figure 3B, S1B - see region where the release size was large enough to avoid drive loss). Another possibility with TADE suppression drive is that the drive might spread too slowly, but this was not a large issue with default migration levels (Figure 3C, S1C). Nevertheless, chasing did substantially affect the performance of TADE suppression drives. With default parameters, suppression without chasing (Figure 3D) was less common than after a period of chasing (Figure 3E). On the other hand, long-term chasing was similarly uncommon (Figure 3F). With ideal parameters, long term chasing was very rare, though there was still usually a period of chasing before suppression occurred (Figure S1D-F).

Fixing the drive release parameters to ensure drive establishment over most of the parameter range, we next assessed the effect of migration rate and low-density growth rate on drive outcomes. When either of these were very low, the drive was unable to establish (Figure 4A), though this was not a problem for the ideal drive (Figure S2A). Drive loss during chasing occurred occasionally, but only when the low-density growth rate was low (Figure 4B). This was likely due to greater stochastic effects under these conditions at low population sizes. When migration was low, the drive was not able to advance quickly, and while some population suppression occurred, some wild-type areas did not experience significant contact with the drive wave of advance by the end of the simulation (Figure 4C). This is a phenomenon specific to TADE drives, not having been seen in much faster zero-threshold suppression drives with the same simulation parameters^42,47^. As migration increased, outcomes shifted to long-term chasing, suppression after chasing, and eventually suppression without chasing (Figure 4D-F). Unlike most other types of drives, the low-density growth rate did not have negative effect on the drive and in fact resulted in slightly more favorable drive outcomes. This is perhaps because the suppression drive wave of advance is boosted by higher low-density growth rate^52^, and higher drive frequency is more helpful to frequency-dependent TADE suppression drives than other zero-threshold drives. Like other types of drives, chasing often ended quickly when migration was higher (Figure 4G), and even during longer-term chasing, the total population size was substantially reduced (Figure 4H), potentially allowing the drive to still have a positive effect. In general, all these patterns held with the ideal drive (Figure S2B-H), with somewhat more favorable drive outcomes across the whole parameter space.

**Figure 4.**
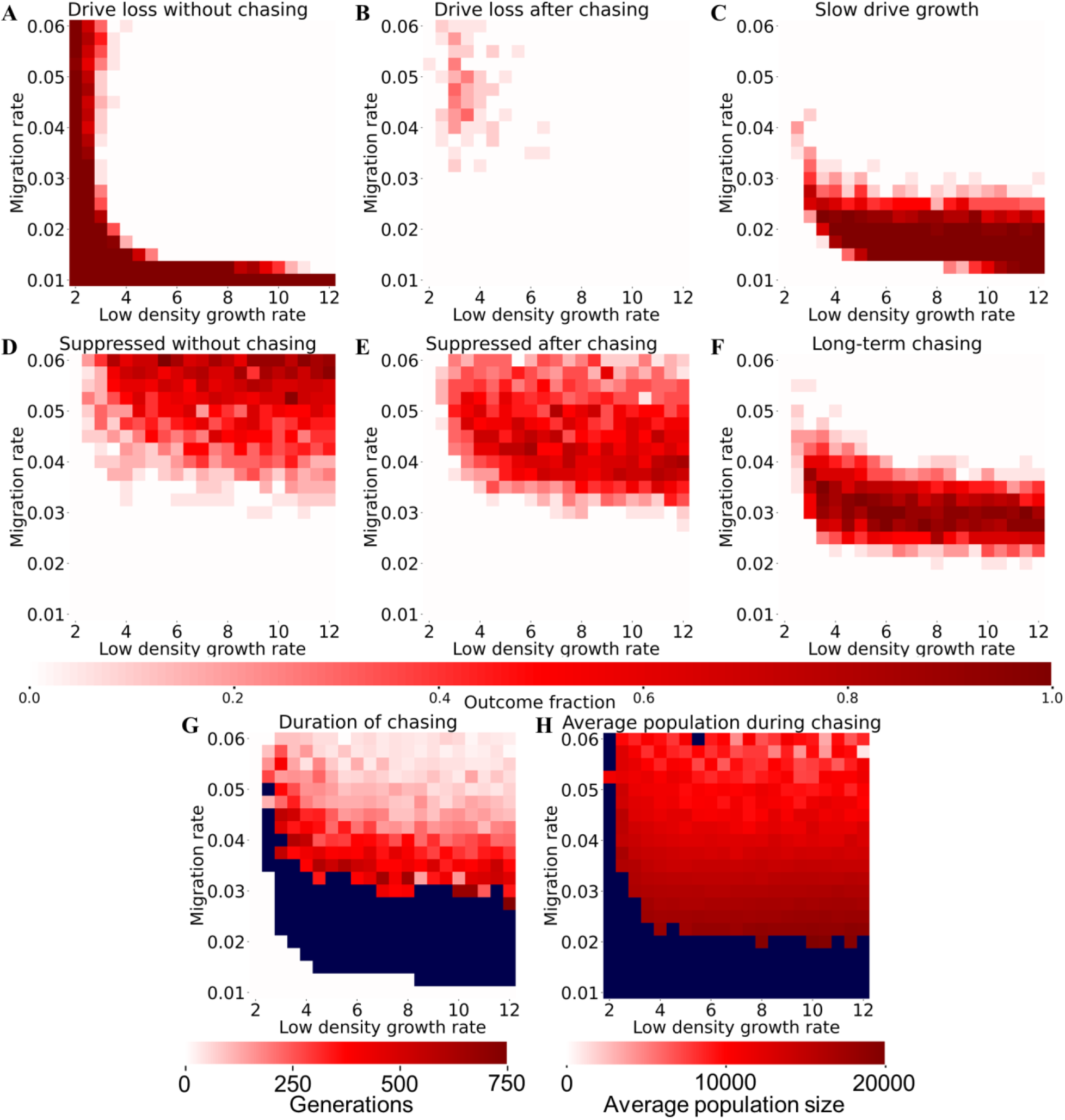
Outcomes with varying ecological parameters. Drive heterozygotes with default parameters and varying migration and low-density growth rate were released into the middle of a spatial population of 50,000 individuals. Outcome rates are displayed for (**A**) drive loss without chasing (usually due to inability of the drive to establish), (**B**) drive loss after a period of chasing, (**C**) simulations in which the drive remained in the population, but spread too slowly to achieve another type of outcome, (**D**) suppression without chasing, (**E**) suppression after a period of chasing, and (**F**) simulations in which chasing was still occurring after 1000 generations had elapsed. Also displayed is (**G**) the average duration of chasing that eventually ended in suppression (blue represents parameter space where the simulation did not end for 1000 generations in any replicate), and (**H**) the average population size during any type of chasing (blue represents parameter space where chasing did not occur). 20 simulations were assessed for each point in the parameter space.

Overall, TADE suppression drive suffered somewhat more from chasing compared to female fertility homing drives and especially TADS suppression drives, but less than both-sex homing suppression drives and Driving Y/X-shredders, consistent with its intermediate population size and expected number of reproducing females in panmictic populations at high drive frequency (see Table S1 and accompanying text).

### Deployment of drives with high embryo cut rates

TADE suppression drive has the potential to advance in space, yet remained confined to a target population. However, construction of ideal or high efficiency TADE drives may not always be possible, or perhaps even more stringent confinement is required for a particular application. In these scenarios, it is worthwhile to consider the properties of TADE suppression drives with much higher embryo cut rates. When 20% or higher, the drive will be unable to sustain a wave of advance. Nevertheless, such drives could still be used to suppress populations with widespread releases. Compared to releases of sterile males and variants of this, continuous releases would certainly be possible and require far smaller numbers. However, panmictic models indicate that a single release of TADE suppression drives might alone be sufficient to eliminate a population if the release size was above the threshold.

To investigate the outcome of a single widespread TADE drive release in our continuous space model, we varied both the embryo cut rate and the release fraction (relative to the normal population size). In general, suppression without chasing was possible over much of the parameter range when the drive was released above its panmictic introduction threshold (Figure 5A), though the release size had to be higher than in panmictic models (to prevent local stochastic failure, which may result in widespread recovery of wild-type). Chasing usually only occurred when the embryo cut rate was low enough to sustain a drive wave of advance (Figure 5B-C). In rare cases, the drive caused local suppression. followed by recolonization by wild-type where the drive remained at low frequency and slowly grew throughout the remainder of the simulation (Figure 5D). Drive loss without chasing occurred mostly when the drive release size was below the threshold (Figure 5E, solid area in the bottom-right), but it could also occur occasionally for higher release fractions (Figure 5E, upper region). In many cases, drive loss occurred after “chasing” was detected (Figure 5F). However, rather than being true chasing, these outcomes usually represented local drive suppression in most areas, but stochastic loss of the drive in some areas where drive frequencies were below the threshold, resulting in a patchy distribution of remaining wild-type followed by wild-type recolonization of the entire arena (Figure 5F). Overall, these results indicate that even worse-performing TADE suppression drives can still succeed in spatial population with widespread releases, albeit requiring higher levels than panmictic populations to ensure success. This conclusion likely would similarly apply to modification drives with a high enough threshold to prevent formation of a drive wave of advance, such as 1-locus 2-drive TARE systems^38^.

**Figure 5.**
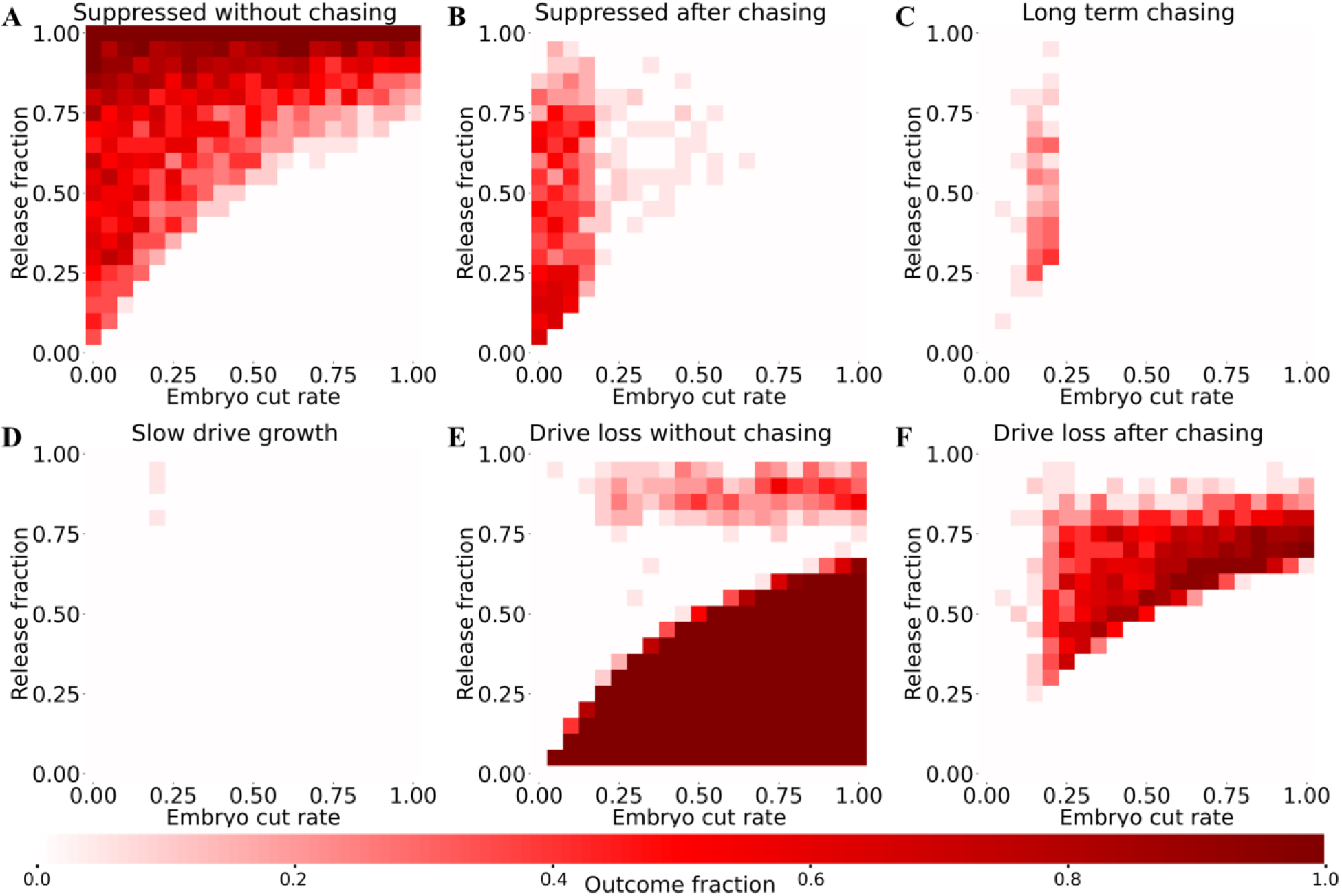
Outcomes with a widespread drive release and varying embryo cut rate. Drive heterozygotes with default parameters were released randomly across an entire spatial population of 50,000 individuals. The embryo cut rate of the drive was varied as well as the release size (as a fraction of the average population size). Outcome rates are displayed for (**A**) suppression without chasing, (**B**) suppression after a period of chasing, (**C**) simulations in which chasing was still occurring after 1000 generations had elapsed, (**D**) simulations in which the drive remained in the population, but spread too slowly to achieve another type of outcome, (**E**) drive loss without chasing (usually due to inability of the drive to establish), and (**F**) drive loss after a period of chasing. 20 simulations were assessed for each point in the parameter space.

## Discussion

Confinement to a target population may be an essential characteristic of gene drives. In some cases, this may be due to sociopolitical considerations, and in other scenarios, it may be essential from a conservation standpoint to avoid suppression of a species in its native range. However, while several available options exist for confined modification drives^15–17,19,37,38,41,54–57^, there are few such designs for suppression systems. Here, we assessed the TADE suppression drive and found that it could be a promising tool for confined suppression of target populations.

In panmictic populations, TADE suppression is a “regional” drive that lacks an introduction threshold in idealized form but where even a small fitness cost or embryo cut rate results in a nonzero threshold^37,38^. Thus, even a somewhat imperfect drive would be expected to spread successfully after a fairly small release in a panmictic population. We found that this was not the case in spatially continuous scenarios. TADE suppression drive was only able to establish in the population and start spreading if released at a moderate frequency over a moderate area. Such dynamics are more similar to “local” drives that have nonzero thresholds in panmictic models even in idealized form. Thus, even a nearly ideal TADE suppression drive could potentially be considered a local drive in populations with a high degree of spatial structure. The reasons for this are likely complex and could potentially be investigated in a follow-up study. However, this phenomenon of greater confinement may be due to the difficulty in forming a drive wave of advance with a frequency-depending drive in situations where the drive is suppressing the population at one side of the wave^52,58^. In continuous space populations, a modification drive can normally advance into a wild-type region if its panmictic introduction threshold is under 50% (or at least fairly close to this level, depending on model detail)^41^. This is not the case for TADE suppression drive. Thus, if an initial drive release is intended to spread through a population after a release in just one part of the population, the embryo cut rate must be sufficiently low. TADE suppression drives with higher embryo cut rates would require widespread or even continuous releases, though such releases would still be far smaller than those involving sterile insect technique-based suppression, even if the sterile males lack any substantial fitness costs. Such TADE drives could be highly confined when the embryo cut rate is over 20% (or lower if the drive had other modest imperfections), even though their panmictic population introduction threshold at this level is well below 50%.

For TADE suppression drives designed to spread through and suppress populations after a single sufficiently large but local release, our modeling indicated that some difficulties might be experienced by the drive due to the chasing effect. This has been shown to potentially be an issue for even powerful global drives with no frequency dependence^42^. However, though TADE suppression drive spreads slower than other previously studied suppression types and requires much higher release fractions to establish and begin suppression, it was not substantially more vulnerable to chasing. This was similar to our previous results with haplodiploid/X-linked homing suppression drives^47^, with the slower suppression of TADE and haplodiploid homing drives potentially allowing them to diffuse more before the population declines, which inhibits chasing (and thus compensates for their reduced overall efficiency). Slower W-shredders had a similar advantage over X-shredders in another modeling study^49^. This suggests that a suitably efficient TADE suppression drive could well be successful in eliminating a target population, though the initial effort needed for the drive to establish is certainly larger than any homing drive. However, high drive efficiency may, in fact, be easier to achieve in a TADE suppression drive than in homing drives because either end-joining or homology-directed repair in the germline could form the desired disrupted allele, while only homology-directed repair in homing drives would provide the desired outcome and contribute to drive success.

While TADE suppression drives are promising, they have not yet been experimentally proven. To construct such a drive, identification of a suitable haplolethal target gene is required, as well as a female fertility gene for drive insertion (though such a drive could also be inserted at the target gene site and target the female fertility gene with additional gRNAs^37^). Presumably, candidates for these genes could be obtained even in less well-studied species by finding homologs of genes in well-characterized species. Additional requirements involve promoters for regulation of drive activity, allowing high cut rates in the germline and avoiding somatic expression, at least in tissues where the haplolethal gene is most essential. These requirements have been met in flies with the *nanos* promoter^22,24^, and some promoters in *Anopheles* may be similarly suitable such as *zpg* and *nanos*^29,32,59^, though these tend to have more somatic expression that may be unacceptably high. Low embryo cut rates would also likely be desired, a characteristic met by the mosquito systems but not yet in fly drives. However, even if all the components for building a drive exist in principle, engineering variants in haplolethal genes may prove difficult because even if the drive was successfully inserted, cutting and end-joining repair at the homologous chromosome’s haplolethal gene may render the cell or organism nonviable. Recoding of the haplolethal gene for efficient rescue could also prove difficult due to the precise requirements for regulating expression of such genes. Nevertheless, both of these issues have been overcome in a homing rescue drive that targeted a haplolethal gene^27^, indicating that TADE suppression drives could be realistically engineered as well.

Aside from TADE suppression drives, other possible options exist for confined suppression. One option is to simply use a homing suppression drive to target alleles that are fixed in the target population, but not found in significant quantity in non-target populations (or at least not fixed if temporary partial suppression and some loss of genetic diversity can be tolerated)^60,61^. While such a drive could be effective, finding a set of target sites that would allow for sufficient gRNA multiplexing to avoid resistance may be a major issue^62^. Another promising type of confined suppression drive is the tethered drive in which a confined modification drive would provide the Cas9 or other components needed by a homing suppression drive^63^. This system was demonstrated successfully in flies^64^. However, even if homing drives avoid resistance alleles, they may fail to suppress populations if the rate of drive conversion/germline homology-directed repair is too low to support a high genetic load^20^. Because any type of repair allows for high genetic load in TADE suppression drives, this could make it a more favored option compared to both of these confined suppression systems in species where high germline cut rates can be obtained but not high rates of homology-directed repair.

With invasive species becoming a more important issue each year due to increasing global trade and climate change, there is a great need for powerful but confined control options that avoid off-target effects on valued species. TADE suppression drive has the promise to meet this requirement. Future studies should thus develop methods for manipulation of haplolethal genes, allowing demonstration of TADE suppression drive in model organisms and eventually pest species of interest. Due to the complexities of gene drive deployment, increasingly realistic modeling is also essential for specific applications and scenarios to ensure that available drives behave in a suitable manner, ensuring both successful population suppression and confinement.

## Acknowledgements

Thanks to Samuel E. Champer and Isabel K. Kim for assistance with SLiM programming and to the High-Performance Computing Platform of the Center for Life Science at Peking University for assistance with cluster-based data collection. This study was supported by laboratory startup funds from Peking University and the NSFC Overseas Youth Fund.

## Supplemental Information

### Supplemental Results

**Figure S1.**
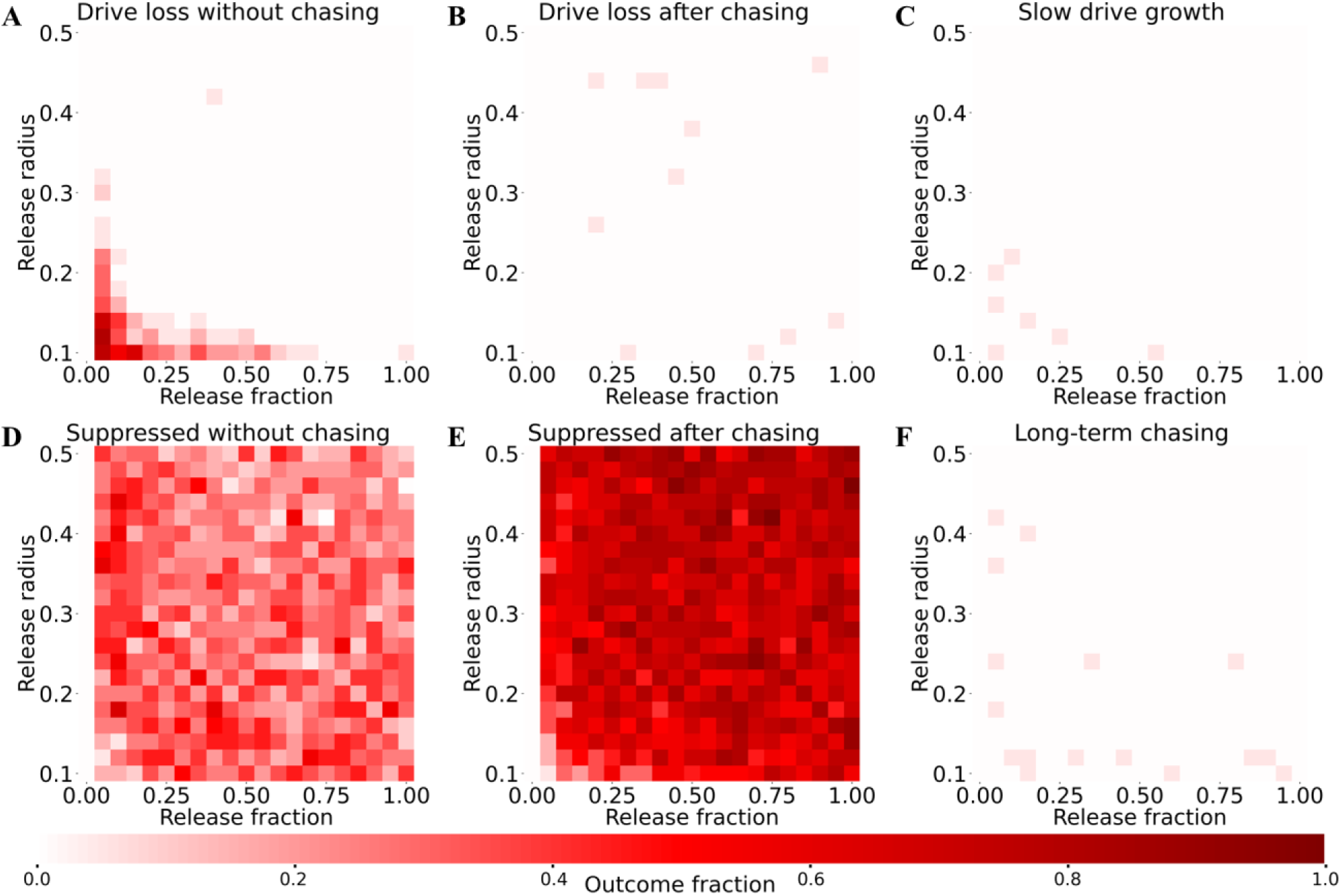
Outcomes with varying release parameters for an ideal drive. Drive heterozygotes with ideal performance were released into the middle of a spatial population of 50,000 individuals. The release radius and the fraction of drive individuals released (proportional to the average population in the release area) were varied. Outcome rates are displayed for (**A**) drive loss without chasing (usually due to inability of the drive to establish), (**B**) drive loss after a period of chasing, (**C**) simulations in which the drive remained in the population, but spread too slowly to achieve another type of outcome, (**D**) suppression without chasing, (**E**) suppression after a period of chasing, and (**F**) simulations in which chasing was still occurring after 1000 generations had elapsed. 20 simulations were assessed for each point in the parameter space.

**Figure S2.**
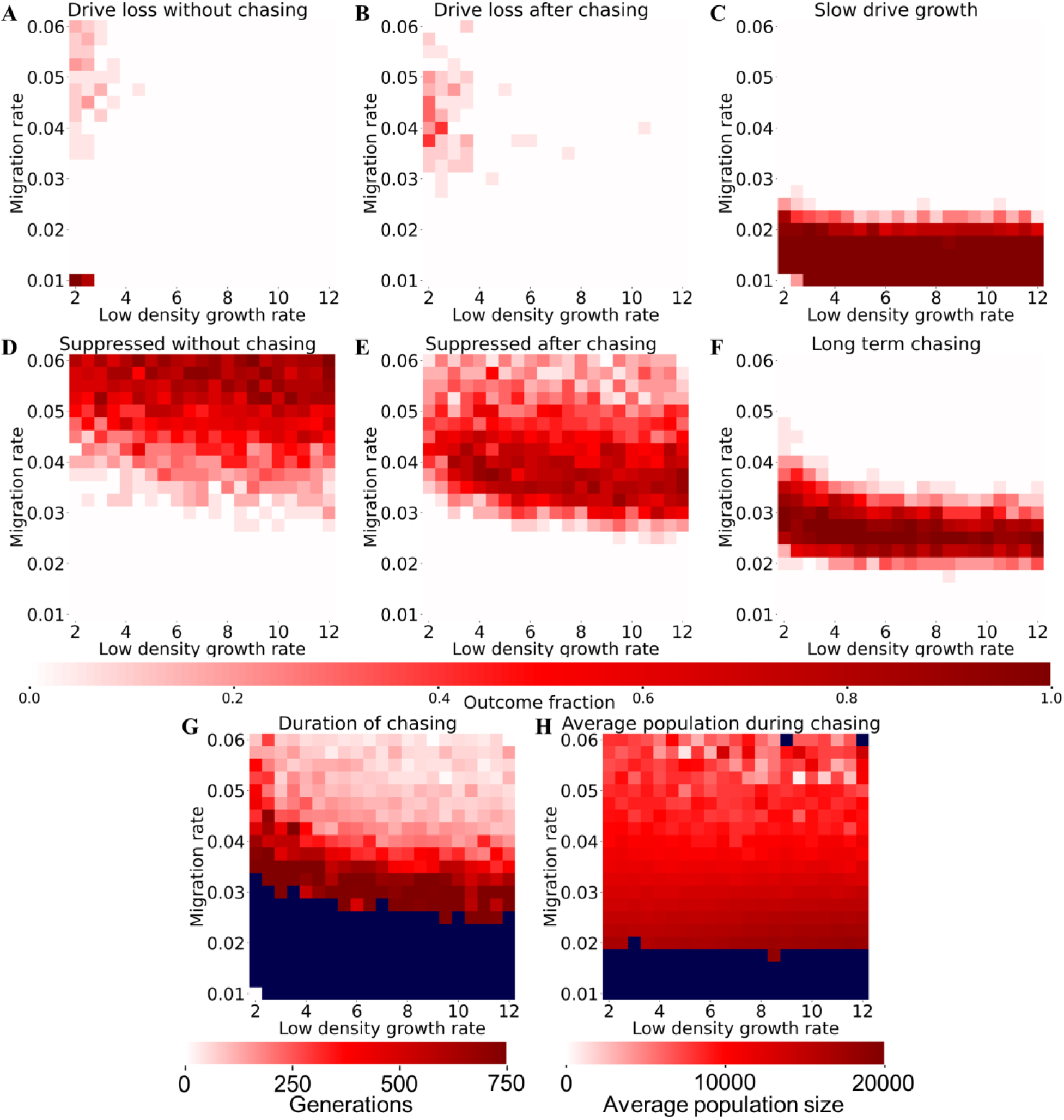
Outcomes with varying ecological parameters for an ideal drive. Drive heterozygotes with ideal performance and varying migration and low-density growth rate were released into the middle of a spatial population of 50,000 individuals. Outcome rates are displayed for (**A**) drive loss without chasing (usually due to inability of the drive to establish), (**B**) drive loss after a period of chasing, (**C**) simulations in which the drive remained in the population, but spread too slowly to achieve another type of outcome, (**D**) suppression without chasing, (**E**) suppression after a period of chasing, and (**F**) simulations in which chasing was still occurring after 1000 generations had elapsed. Also displayed is (**G**) the average duration of chasing that eventually ended in suppression (blue represents parameter space where the simulation did not end for 1000 generations in any replicate), and (**H**) the average population size during any type of chasing (blue represents parameter space where chasing did not occur). 20 simulations were assessed for each point in the parameter space.

**Table S1.** Population characteristics at 90% drive frequency correlate with chasing. The table shows population characteristics for the generation in which the drive is closest to 90% allele frequency in a panmictic population of 100,000 (using default parameters for each). In a spatially continuous model, this approximately corresponds to a part of the drive wave of advance furthest from the wild-type population and where population elimination is imminent. New wild-type individuals generated in this area can potentially move into the adjacent empty region and begin chasing. Higher levels of stochasticity tend to support this process. Thus, the both-sex fertility homing drive with low numbers of reproducing females is particularly prone to chasing, while the TADS suppression drive can generally avoid chasing. The Driving Y has an intermediate number, but overall population density is high, resulting in fewer offspring and stronger stochastic effects for the reproducing individuals. The haplodiploid suppression drive is intermediate between the autosomal female fertility homing and TADS suppression drive, but it is similarly (or slightly more) vulnerable to chasing as the autosomal female fertility homing drive, perhaps due to other factors that cannot be captured in this analysis such as lower overall drive power or genetic load. The TADE suppression drive is similar to the female fertility drive in this table and is perhaps slightly more vulnerable to chasing after the drive is established. It is slower to increase, but it can retain a high genetic load^37,38^.

|  | TADE suppression | Haplodiploid female fertility homing | Female fertility homing | Both-sex fertility homing | Driving Y (X-shredder) | TADS Suppression |
| --- | --- | --- | --- | --- | --- | --- |
| Population size | 872 | 3404 | 1363 | 865 | 10969 | 10999 |
| Fertile females | 11% | 10% | 7.9% | 8.1% | 9.1% | 50% |
| Sterile females | 39% | 40% | 43% | 41% | N/A | N/A |
| Fertile males | 50% | 50% | 49% | 8.8% | 8.2% | 8.3% |
| Sterile males | N/A | N/A | N/A | 42% | 83%* | 42% |
| **Expected reproducing females** | **93** | **345** | **114** | **12** | **103\*\*** | **934** |
| **Long-term chasing rate** | **medium-high** | **medium** | **medium** | **very high** | **high** | **low** |

Values are the average over 20 simulations. Except for the TADE suppression drive, data is from previous studies<sup>42,47</sup>. \*Drive-carrying males. \*\*Includes only females expected to mate with wild-type males.

## Notes

https://github.com/jchamper/ChamperLab/tree/main/TADE-Suppression-Modeling

### Competing Interest Statement

The authors have declared no competing interest.

